# Wireless AI-Powered IoT Sensors for Laboratory Mice Behavior Recognition

**DOI:** 10.1101/2020.07.23.217190

**Authors:** Meng Chen, Yifan Liu, John Chung Tam, Ho-yin Chan, Xinyue Li, Chishing Chan, Wen J. Li

## Abstract

According to the U.S. Department of Agriculture in 2018, there are more than 100 million animals used in research, education, and testing per year. Of the laboratory animals used for research, 95 percent are mice and rats as reported by the Foundation for Biomedical Research (FBR). We present here our work in developing wireless Artificial Intelligent (AI)-powered IoT Sensors (AIIS) for laboratory mice motion recognition utilizing embedded micro-inertial measurement units (uIMUs). Based on the AIIS, we have demonstrated a small-animal motion tracking and recognition system that could recognize 5 common behaviors of mice in cages with accuracy of ~76.23%. The key advantage of this AIIS-based system is to enable high throughput behavioral monitoring of multiple to a large group of laboratory animals, in contrast to traditional video tracking systems that usually track only single or a few animals at a time. The system collects motion data (i.e., three axes linear accelerations and three axes angular velocities) from the IoT sensors attached to different mice, and classifies these data into different behaviors using machine learning algorithms. One of the challenging problems for data analysis is that the distribution of behavior samples is extremely imbalanced. Behaviors such as *sleeping* and *walking* dominate the entire sample set from different mice. However, machine learning algorithms often require a balanced sample set to achieve optimal performance. Thus, several methods are proposed to solve the imbalanced sample problem. Data processing methods for data segmentation, feature extraction, feature selection, imbalanced learning, and machine learning are explored to process motion data including *sleeping, walking, rearing, digging, shaking, grooming, drinking and scratching*. For example, by tuning the parameters of a machine-learning algorithm (i.e., Support Vector Machine (SVM)), the average accuracy of classifying five behaviors (i.e., sleeping, walking, rearing, digging and shaking) is 48.07% before solving the imbalance sample issue. To address this problem, an iteration of sample and feature selection is applied to improve the SVM performance. A combination of oversampling and undersampling is used to handle imbalanced classes, and feature selection provides the optimal number of features. The accuracy increases from 48.07% to 76.23% when the optimized combination is used. We further obtained an average accuracy of 86.46% by removing *shaking*, which is proved to have a negative effect on the overall performance, out of these five behaviors. Furthermore, we were able to classify less frequent behaviors including *rearing, digging, grooming, drinking* and *scratching* at an average accuracy of 96.35%.

## 1. Introduction

Animal studies are essential in drug discovery [1], biomedical engineering, and biological research [2]. For example, animal behavior is often used to learn how certain genes are involved in learning and memory functions [3] or to evaluate the effectiveness of candidate drugs prior to human trials [4]. In addition, detailed behavior analysis of individual animal allows the early detection of abnormal behaviors [5], which results in better animal care [6]. Until now, behavioral monitoring of lab animals is based almost solely on video recording [7-10], which is extremely time-consuming and labor-intensive because each camera can only record and analyze one or maximally a few animals at a time. In addition to the high cost, such inefficiency leads to limited sample size, and hence poor reliability and consistency of animal studies. Although commercial video-based tracking systems (e.g., EthoVision XT by Noldus and Topscan by CleverSys) that track animal behaviors in real-time are available, their accuracy can reach only 70% [11] with limited number of tracked animals due to the limited field of view and camera resolution. These problems hinder the progress of science, medicine, and public health. Thus, a low-cost solution using Artificial Intelligence (AI)-powered animal motion tracking system is proposed to enable both individual and multiple behavioral monitoring of lab animals. We are developing motion tracking devices in order to ultimately achieve the following two functions: (1) measure different types of behaviors, such as walking, circling, rearing, and social interaction, which provide important phenotypic readout for biomedical and drug discovery (especially neuroscience/neurology), and (2) simultaneously monitor a large number (hundreds) of animals to address the low efficiency and poor data quality of existing behavioral monitoring systems. To the best of our knowledge, no system that delivers similar functions exists at present. Although several commercial products are capable of measuring other physiological data (e.g. Electroencephalography (EEG) [12-14] and blood pressure [15-17]), the TSE and EMKA systems can only measure the presence or absence of activities, but cannot differentiate different types of behaviors. Thus, these systems cannot monitor behavior. More importantly, these systems are unsuitable for large-scale studies because they require highly complicated and time-consuming (3–4 hours per animal) surgical procedure to implant the device inside the animals. Moreover, highly trained specialists are needed for the procedure. By contrast, our target device is designed to be small enough to be delivered by subdermal injection. Each injection takes no more than five seconds and no specially trained technicians are required. Thus, the device is applicable for a large number of animals.

Acceleration based activity/behavior/motion recognition has recently drawn attention because of the success of the industry’s first MEMS 3-axis inertial sensor in 2012 [18]. The application starts from dedicated accelerometers [19] to embedded accelerometers [20] in mobile phones, which are used for monitoring human physical activities. The classification accuracy of human behaviors can achieve over 95% for five behaviors including running, standing, biking, sitting, and walking [21]. Machine learning algorithms have commonly been used to classify behaviors such as decision tree [22], K-nearest neighbors [23], Support Vector Machine (SVM) [24], Naive Bayes [25], and Neural Networks [26]. Decision tree and Naïve Bayes are based on statistics, which requires much data to form relative reliable probabilities for each class. K-nearest neighbors only concerns the nearest K points of a new input, which is probably surrounded by points from majority class. Neural Networks tune parameters by iterations of trainings to achieve optimal performance. This model tends to satisfy the classes with the most training data, and thus is not applicable to an imbalanced data set. However, SVM relies on kernel functions that project all instances to a higher dimensional space with the aim of finding a linear decision boundary (i.e., a hyperplane) to group the data. Only the points lying on the boundary affect the performance of the classifier. The distribution of majority of the classes does not have critical effects. Therefore, this method is suitable for an imbalanced data set.

Animal motion tracking is more difficult than that of human because animal behaviors are uncontrollable. The distribution of the collected motion sample is extremely imbalanced. For instance, mice sleep most of the day and this act contributes over 80% of the total motion samples. Unfortunately, machine learning algorithms require balanced training data to achieve their performance. Thus, imbalanced learning was proposed to deal with an imbalance data set, which often occurs when mining data for text categorization to biomedical data analysis [27]. The strategies [28] used to overcome imbalance sample distribution can be categorized into sampling strategy [29], synthetic data generation [30, 31], cost-sensitive learning [32], active learning [33], and kernel-based methods [34]. Sampling strategy [22] develops oversampling and undersampling methods to balance the distribution in the original data set. Synthetic data generation aims to generate more data for the minority classes to regain the balance in the original dataset, such as Synthetic Minority Over-sampling Technique (SMOTE) algorithm [30] and its extension SMOTE-BOOST [31]. Cost-sensitive learning provides a cost coefficient for each class in its machine learning model [32] to reduce misclassification without modifying the original dataset. Active learning [33] can select most relevant samples for training to reduce the computational cost. Kernel-based methods are proposed to solve imbalanced learning by modifying the kernel matrix [34] according to the distribution of classes in training data. In this paper, both sampling strategy and active learning are combined to realize sample selection and generate optimal balanced training data set.

To improve the utilization of the limited motion data collected from animal experiments, noise or irrelevant information (such as irrelevant features) should be removed from training samples. In this work, six features including mean, variation, root mean square, maximum, minimum values, and energy [35] of each axis (i.e., x-, y-, z-directions of acceleration and rotation) are extracted from motion data. A good set of extracted features should well and efficiently represent the input data and is independent from each other. Feature selection (variable elimination) helps in understanding data, reducing computation requirement, reducing the effect of the curse of dimensionality, and improving predictor performances [36]. Feature selection methods can be categorized into filter [37] and wrapper methods [38]. Filter methods select high ranking (i.e., most important) features that are independent of classifiers whereas wrapper methods evaluate features according to classifier performance. Filter methods remove features that are irrelevant to class labels through signal processing methods, and then the selected features are used to test the performance of different classifiers. However, these features may have significant effects in specific classifiers. Given that a classifier is chosen, wrapper methods are more suitable for feature selection.

In this paper, an animal motion tracking system based on the wireless AI-powered IoT sensors is proposed, as shown in Figure 1, to monitor and track small animal motions by analyzing the collected accelerations and angular velocities. We build the wireless AI-powered IoT sensor, named as motion tracker in this project, that is small and light enough for a small animal to avoid negative influence on its motion. By utilizing a low-power wireless transceiver, the tracker can run without wires for a long period. Using the developed platform, several experiments on C57BL/6 mouse were conducted to collect the mice’s motion data, which were classified into five basic behaviors of sleeping, walking, rearing, digging, and shaking. The data processing procedures that include segmentation, feature extraction, and imbalanced learning are discussed.

**Figure 1.**
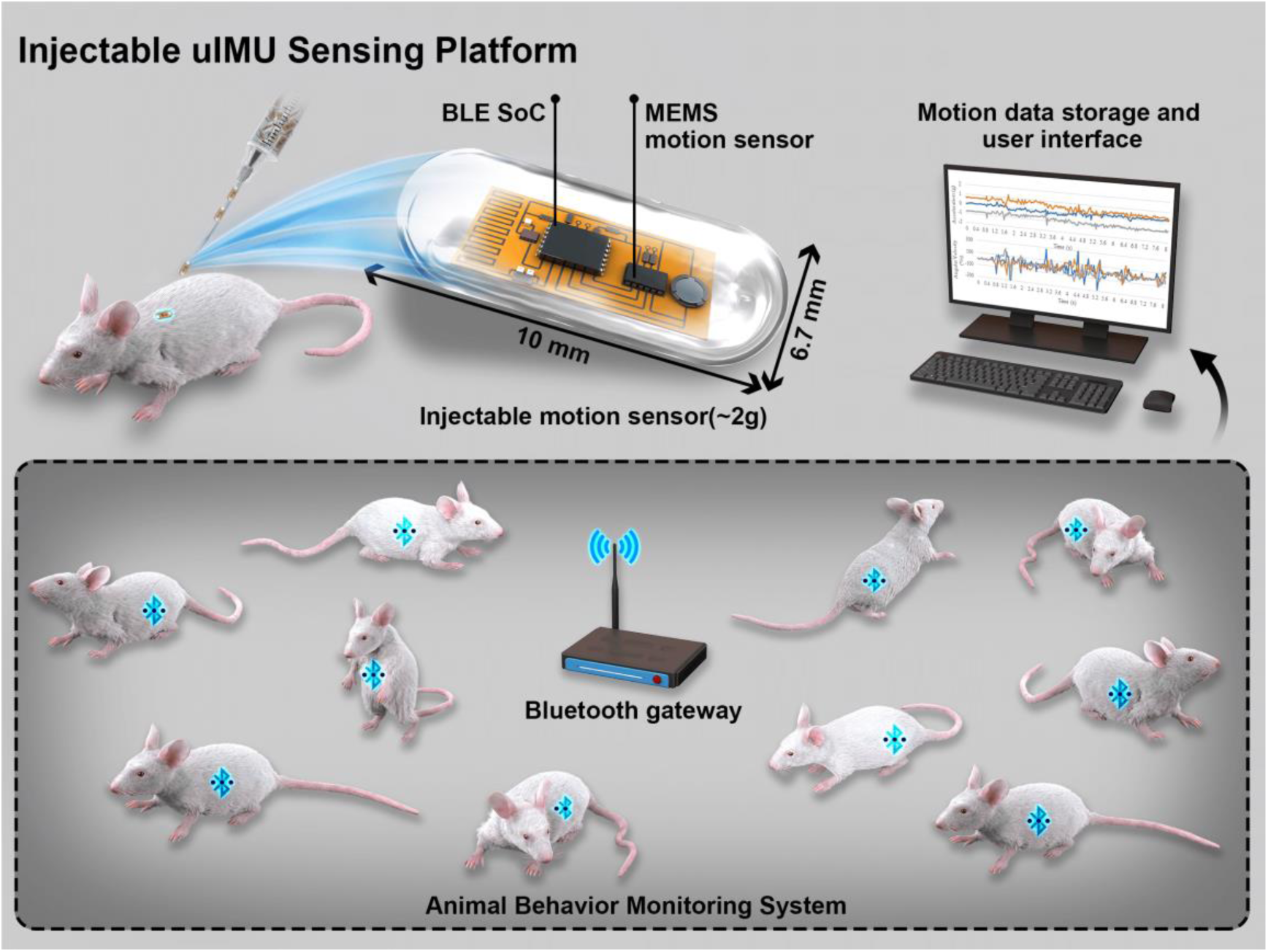
Conceptual illustration of an ***injectable uIMU sensing platform*** for tracking multiple mice behavior recognition

## 2. Artificial Intelligence (AI)-powered animal motion tracking system

### 2.1 Overall system design

Figure 2(a) shows the overall system design. A dedicated motion tracker, as shown in Figure 2(b), is mounted on a mouse’s back to monitor the mouse’s motion. One gateway and multiple trackers form a sensor network to implement multi-target--monitoring. Cameras are used to record synchronized videos of the mice’s motion for behavior labeling. The motion data are then forwarded to a cloud server for storage and further behavior classification.

**Figure 2.**
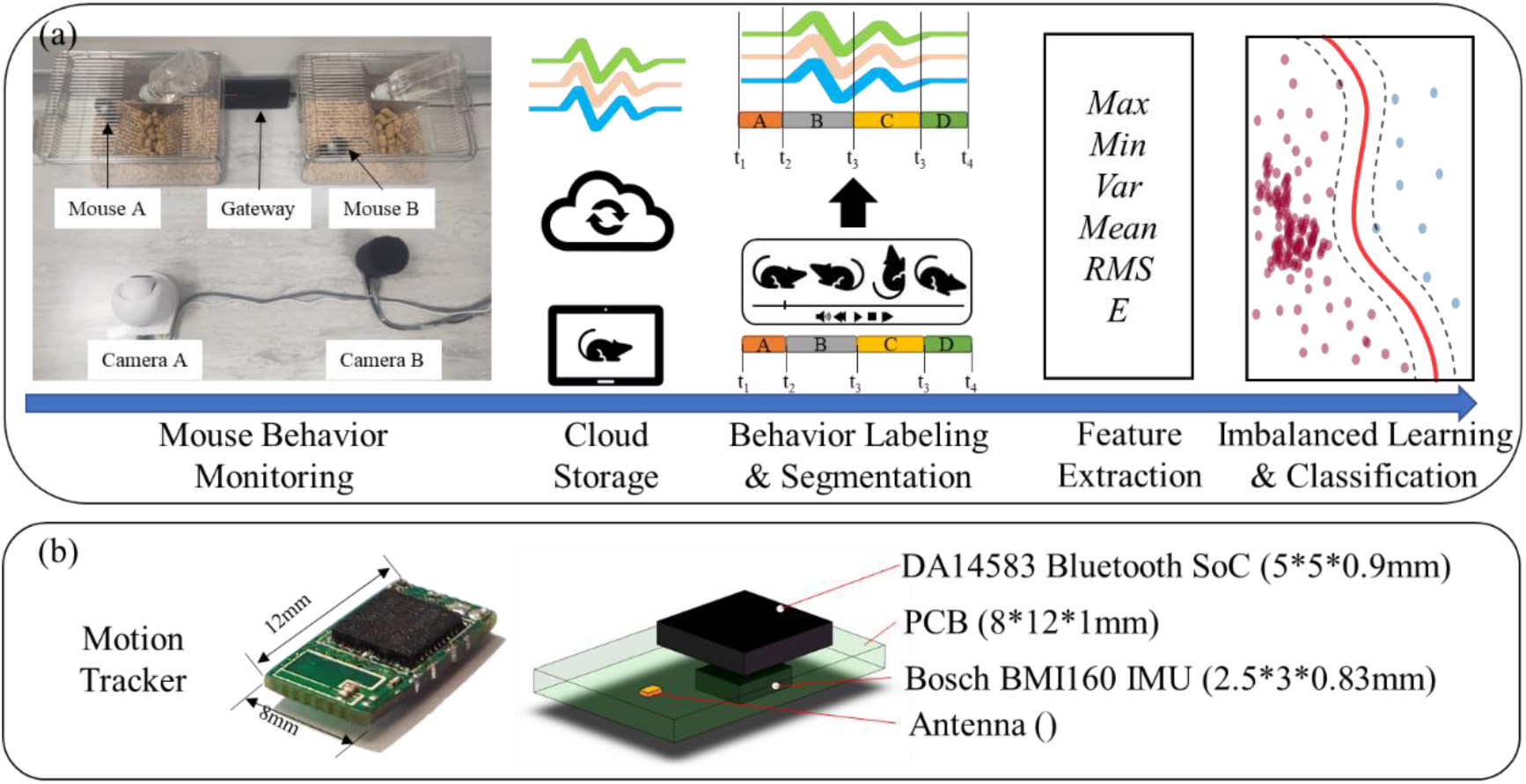
(a) AI-powered motion tracking system; (b) Design of a motion tracker

For hardware part, the motion tracker (Figure 2(b)) is mounted on monitoring targets such as mice whose sizes are less than 50g. Thus, to minimize the size and weight of the sensor, a Bluetooth Soc (Dialog DA14583) is integrated to serve as a microcontroller and a transceiver, and a small low energy IMU chip (Bosch BMI160) is applied to measure both acceleration and angular velocity. The communication between the tracker and a receiver is achieved by Bluetooth Low Energy (BLE) which consumes less energy but has an acceptable data rate and communication range as compared with several IoT protocols, such as Wi-Fi, Bluetooth, and ZigBee, as shown in Table 1.

**Table 1.** IoT Protocols comparison [39]

| IoT Protocols | Max. Nodes per Master | Peak Current Consumption | Range | Data Rate | Topology | Relative Cost |
| --- | --- | --- | --- | --- | --- | --- |
| Wi-Fi: IEEE 802.11b | 32 | ~100 mA | 100 m | 54 Mb/s | Star | Medium |
| Bluetooth | 7 | 30 mA | 10 m | 1 Mb/s | Star | Low |
| ZigBee | 100 | 30 mA | 50 m | 250 kb/s | Star/Mesh | Low |
| Bluetooth Low Energy | 7+ | 15 mA | 100 m+ | 1 Mb/s | Star/Mesh | Low |

In this small sensor network, the receiver acts as a bridge to connect data transmission from the tracker to a cloud server. Thus, both BLE and Wi-Fi are supported. The receiver has sufficient computational capacity for local data processing, such as data aggregation and compression. A smartphone or a single board computer like Raspberry Pi can sufficiently satisfy the requirements of such receiver. In this project, a Nokia X71 Android smartphone is used and continuously forwards the collected motion data to a cloud server. To record mouse behaviors for further labelling process, a video surveillance system that includes a camera and a local NAS (Network Attached Storage) server is set. The camera is placed at the top or on the side of the cage to have a better view of the full cage. The recorded video data have timestamps, which provide synchronization with motion data collected by the tracker.

For the software, the data processing steps include behavior labeling, segmentation, feature extraction, and imbalanced learning. First, behavior labeling is applied manually on the motion data collected by the tracker according to the recorded synchronized videos. The labels split the motion data into pieces, each of which is further segmented by a time window. Each data segment has 36 features that are extracted to form one sample set. Imbalanced learning rebuilds a balanced training data set. Feature selection chooses the optimal combination. The parameters of the two methods are tuned by the performance of an SVM classifier. Finally, the classifier provides a behavior classification for each sample set.

### 2.2 Procedures for motion tracker mounting

Figure 3 shows the procedures for a motion tracker mounted on a mouse (C57BL/6). First, flexible tubing made of elastic rubber was placed around the neck of the mouse to avoid direct contact of the sensor and to act as a ground for sensor mounting. The sensor was wrapped by a piece of parafilm to seal the electronics parts. Then, the mouse was anaesthetized in a chamber by isoflurane, a kind of inhalation anesthetic with relative short onset and recovery time, as shown in Figure 3(b). Under total anesthesia, the mouse was transferred and masked by an isoflurane anesthesia machine with a tube, as shown in Figure 3(c). Next, the flexible tubing was wrapped tightly around the upper part of the mouse, which was between the front and back legs. Then, our tracker was attached at the back of the mouse by tape, its actual location is depicted in Figure 3(e). The tracker was oriented such that the x-axis of the sensor should be along with the mouse’s spine. The y-axis should be parallel with the line connecting the two ears of the mouse.

**Figure 3.**
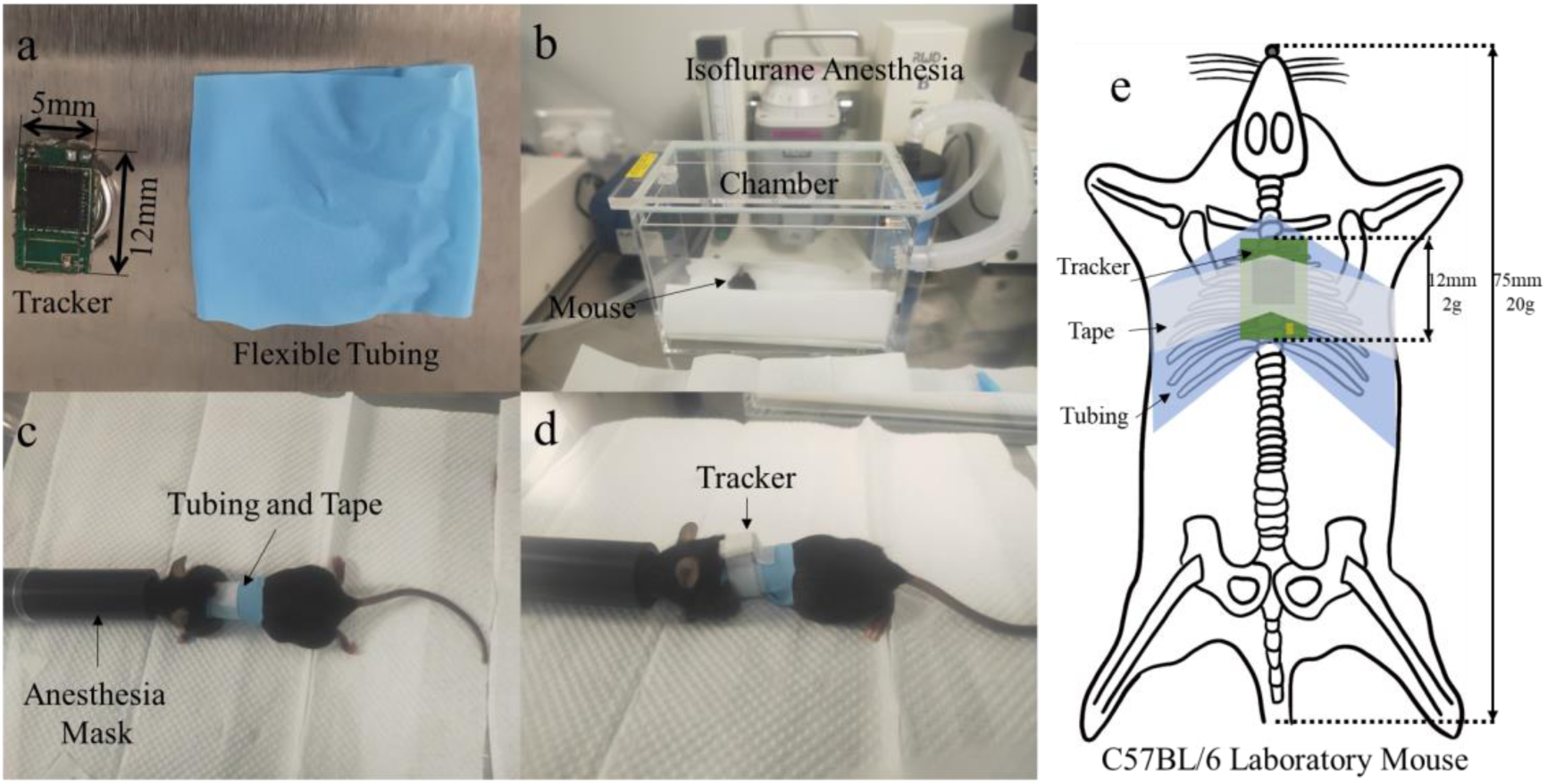
Procedures of mounting a tracker on a mouse. (a) prepare the tracker and flexible tubing; (b) anesthetize the mouse; (c) wrap the tubing around the mouse; (d) mount the tracker; and (e) tracker location

### 2.3 Data collection of multiple animals

The setup of multi-animal experiment is shown in Figure 4. There are two reasons why four cages were used instead of a single cage. First, the tracking devices were mounted at the back of each mouse, and they would have a high chance of being taken down among each other if they were placed in the same cage. Second, manual labeling of behaviors is difficult when four mice were placed in the same cage. Those mice may block the view of cameras. This is also the main advantage of our developed system which relies only on sensor data for behavioral monitoring and solves the camera blockage problem.

**Figure 4.**
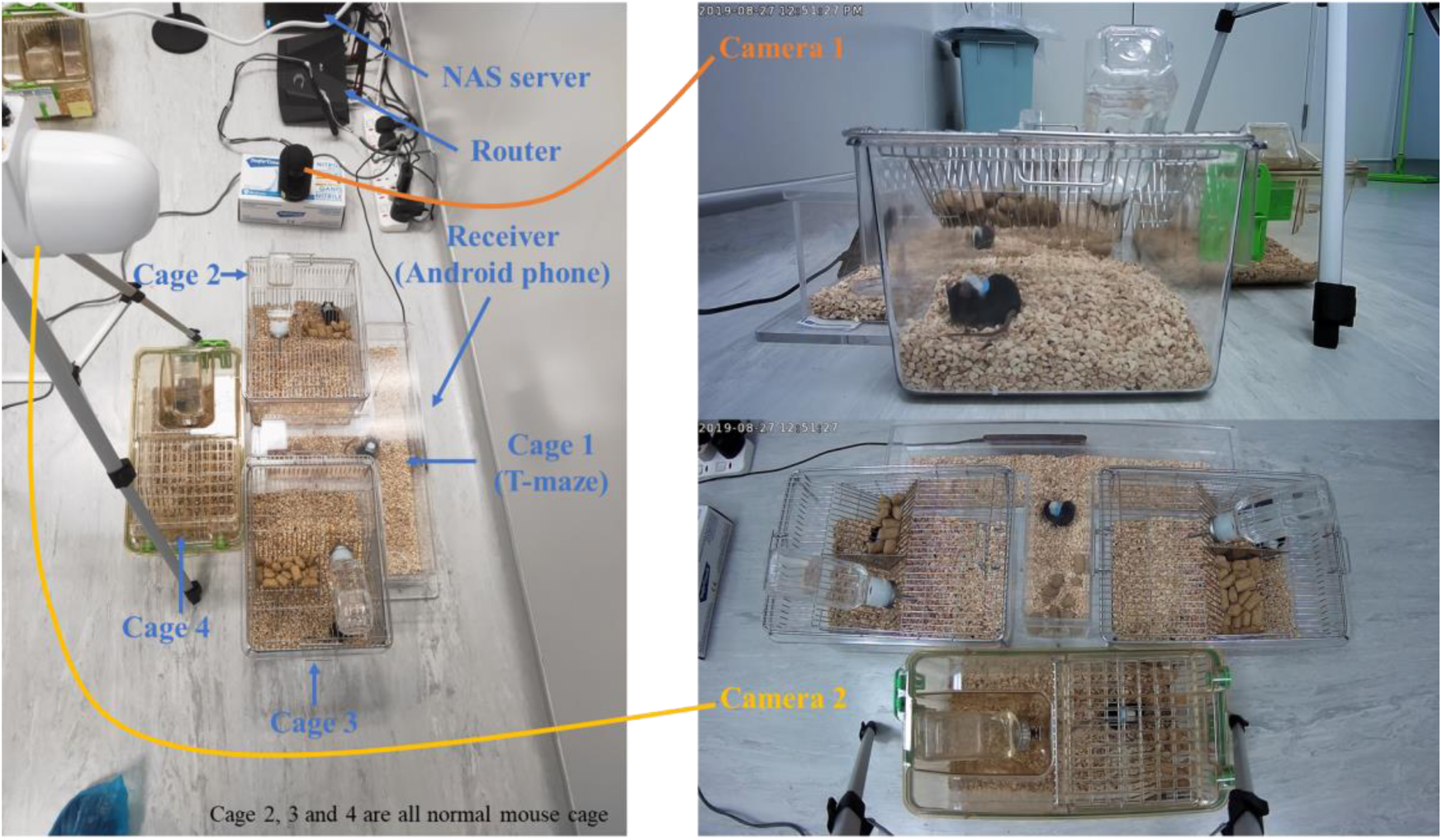
Setup of multiple mouse experiment (Left) and views of video captured by the two cameras (Right)

In our experiment, the recorded videos were stored in a local NAS server. An Android smart phone (Nokia X71) was placed beside a cage as a receiver that connects all the four tracking devices. The phone collects all motion data (i.e., acceleration and angular velocity) and forwards them to a remote server. Figure 5 shows examples of motion data collected from all the four tracking devices. It is shown that the developed tracking platform can simultaneously monitor multiple animals.

**Figure 5.**
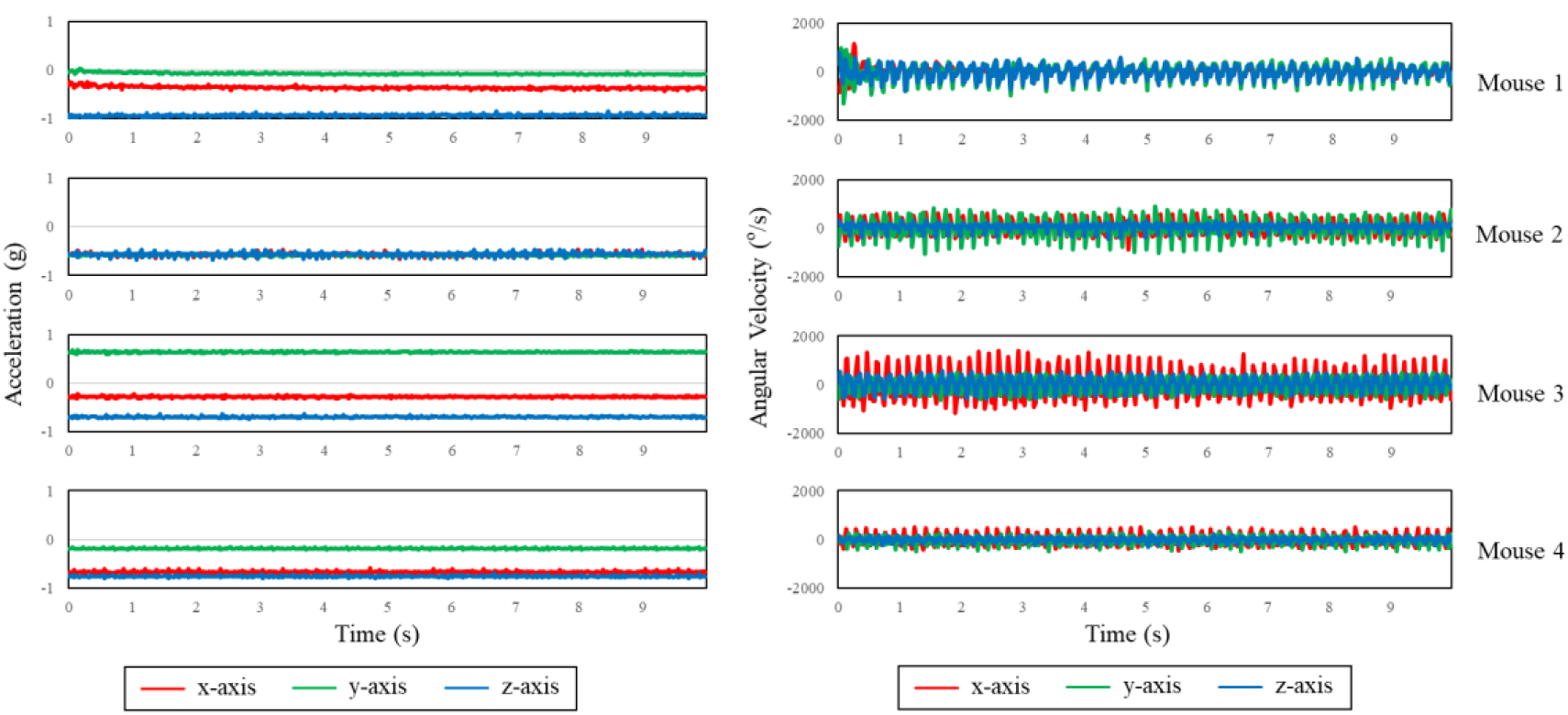
Motion data (acceleration and angular velocity) of multiple animal experiment

## 3. Data Analysis Methodology

### 3.1 Fixed-time-window Segmentation

Segmentation splits the raw motion data into pieces according to their behavior labels. These labeled motion data required for machine learning training and testing are prepared by referencing the corresponding video data. As discussed earlier, a camera records the motion video during the experiment. The videos are synchronized with the motion data according to the time stamp of each sensor reading. The start and end times of a specific behavior label are recorded manually by studying the videos. These time stamps are used to segment and group the motion data into different clusters (i.e., labelled motion data). Then, the labelled motion data are further segmented using fixed-time windows (e.g., 1s, 2s, and so on). Each data segment comprises one sample. The selection of window segment depends on the shortest duration of labeled behaviors.

### 3.2 Feature extraction and selection

Each data segment contains information of three-axis accelerations and three-axis angular velocities. For each axis, six time-domain features are extracted that include mean, variance, maximum, minimum, root mean square, and energy, as shown in Table 2. Thus, 36 features are derived for each data segment, and a feature set is created. To determine the most problem-relevant features, a Sequential Feature Selection (SFS) algorithm is used to reduce an initial *d*-dimensional to a *k*-dimensional feature subspace (*k* < *d*) using greedy search algorithm. This algorithm in turn removes or adds one feature at a time based on classifier’s performance until a feature subset of the desired size *k* is reached [40]. The selected feature subset is much smaller than the original feature set. Thus, computational efficiency can be improved. Furthermore, irrelevant features are removed to reduce the generalization error of the model that is very important as discussed in next section.

**Table 2.**
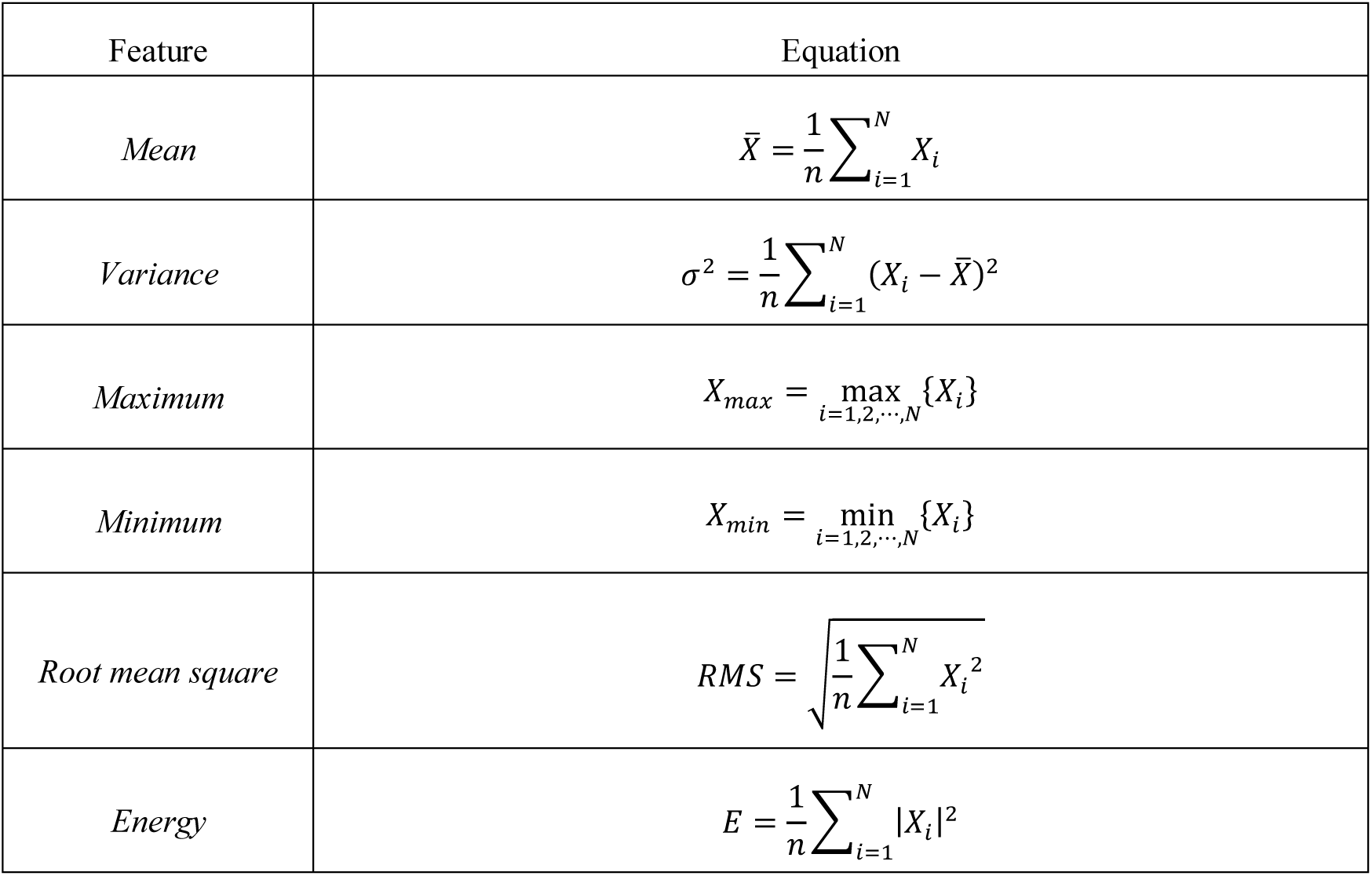
Six time-domain features extracted from each axis

### 3.3 Imbalanced learning

An imbalanced data set means the number of samples in one classification is far more than that of samples in other classifications. Most standard algorithms assume or expect balanced class distributions or equal misclassification costs. During the training process, the loss or error changes once for one training sample. A balanced training set provides equivalent contribution on loss or error for each class. However, for an imbalanced training set, the loss or error changes frequently for majority classes. Then, the model tends to lean toward the majority classes. Therefore, when presented with complex imbalanced data sets, these algorithms fail to properly represent the distributive characteristics of the data and provide unfavorable accuracies across data classes [41]. Random sampling is a typical method to solve the imbalance by adjusting the class distribution of a data set. Two kinds of random sampling methods can be applied: oversampling and undersampling. Oversampling is achieved by adding further samples to minority classes, which can have as many samples as the majority classes. SMOTE [30] is the most common technique of oversampling and proved the feasibility in various areas. The method artificially synthesizes new samples by calculating the similarity of feature spaces based on samples in the minority class. The procedure is introduced as below.

1. Find *k* nearest neighbors for *x*_*i*_, where *x*_*i*_ is a sample of the minor class.
2. For each neighbor 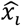, generate one new sample *x*_*new*_ according to:

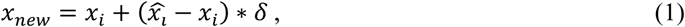

where δ ∈ [0,1] is a random number.
3. Add the new sample back to the data set.

Undersampling is achieved by removing several samples from majority class to keep balance with other classes. ClusterCentroids achieves undersampling by generating centroids using clustering methods. *N* clusters are generated from the majority class using *k*-means algorithm. The *N* centroids of the *N* clusters are the new samples of oversampling the majority class to *N* samples.

1. Randomly initialize *N* cluster centroids *μ*_1_, *μ*_2_, ⋯, *μ*_*N*_.
2. For each cluster centroid *μ*_*i*_, find *k* nearest points.
3. Relocate the cluster centroid *μ*_*i*_ by calculating the mean of the cluster.
4. Repeat step 3 until:

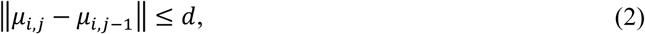

where *μ*_*i,j*_ is the *j*th cluster centroid and *d* is a threshold.
5. The *N* cluster centroids *μ*_1_, *μ*_2_, ⋯, *μ*_*N*_ are the generated samples.

In this study, a sample selection is used, which is the combination of both the ClusterCentroids and SMOTE to generate a certain size of samples. To convert a training data set from an imbalanced to a balanced one, a suitable number of samples for each class is determined by classifier’s performance. This number is usually laid between that of samples in a minority class and that in a majority class. Thus, a search process in the range is applied to find the suitable number with the highest classification accuracy. To avoid the overfitting problem, we use the five-folder cross validation during the training process. The operation of both undersampling and oversampling must be on the whole dataset (i.e., all classes), and thus the ClusterCentroids is first applied to reduce the number of samples of majority class, and then SMOTE is used to increase the number of samples of all classes to a suitable size.

### 3.4 Classification

The SVM presents numerous advantages over other machine learning algorithms. The SVM is a type of machine learning method that works well with small dataset and different from statistical ones that rely on possibility density prediction and law of large numbers. The SVM decision function only relates to support vectors.

The SVM is designed to solve the two-class problem by finding a hyperplane to divide the data set into two classes [42]. The principle of dividing is to maximize the margin between two classes, and finally solves the convex quadratic programming problem. If a data set is linearly separable, hard margin method is used to train a linearly separable SVM. If a data set is approximately linear separable, a soft margin is used to train a linear SVM. If a dataset is linearly inseparable, kernel and soft margin are used to train a nonlinear SVM. In this study, a radial basis function kernel was used.

## 4. Experimental Result

### 4.1 Behavior Motion Visualization

Five of the most frequent behaviors of a mouse, namely, *sleeping, walking, rearing, digging*, and *shaking* are chosen in this study. By studying the recorded videos during experiments, these five behaviors are identified and labeled manually. The starting time, ending time, and behavior classification of each detected behavior are then recorded in a list. Given that the video and motion data are captured simultaneously, finding the detected behaviors in mouse’s motion data can simply done by matching the corresponding time stamps between the two datasets. According to the starting time, ending time, and behavior classification, the motion data are extracted from the entire motion data labeled by class.

The labeled motion data are further split into pieces by a two-second time window. The determination of using two seconds will be explained later. To visualize the difference of five behaviors from the perspective of motion data, eight-second labeled motion data from each classification are compared, as shown in Figure 6. Sample videos of different behaviors are attached in the supplementary section. The motion data of these five behaviors are studied and listed in Table 3.

**Table 3.** Summary of motion change in sleeping, walking, rearing, digging, and shaking

| Behaviors | *Acceleration* |  |  | *Angular Velocity* |  |  |
| --- | --- | --- | --- | --- | --- | --- |
| | $a_x$ | $a_y$ | $a_z$ | $g_x$ | $g_y$ | $g_z$ |
| **Sleeping** | No change | No change | No change | No change | No change | No change |
| **Walking** | Small<br>Periodical | Small<br>Periodical | Small<br>Periodical | Large | Small | Small |
| **Rearing** | Medium | Large | Large | Large | Large | Large |
| **Digging** | Large | Large | Large | Large | Large | Large |
| **Shaking** | Medium | Medium | Medium | Large | Large | Large |

**Figure 6.**
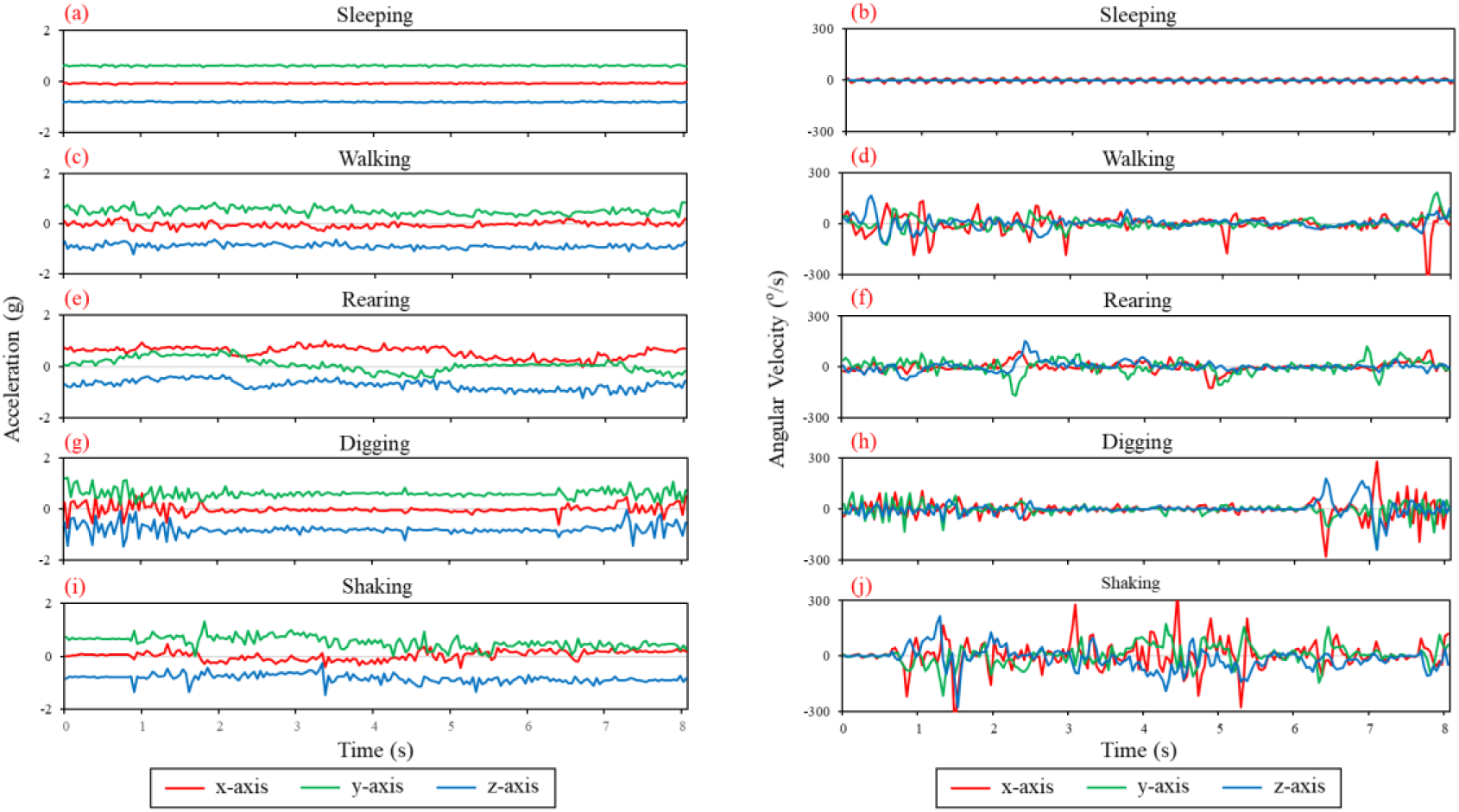
Three-axis acceleration and angular velocity of sleeping, walking, rearing, digging, and shaking (Duration: 8 second)

Among the five behaviors, sleeping is the most frequent because of its long and continuous period. The mouse keeps still except for breathing, which causes small variations on the accelerometer and gyroscope readings. Figures 4(a) and 4(b) show the eight-second acceleration and angular velocity of a sleeping mouse, respectively. As expected, the acceleration and angular velocity remain stable while the mouse is sleeping. Walking is a transition behavior. The mouse rarely continues walking because the cage is so small and does not have enough space. Before and after walking, the mouse always performs other behaviors. During walking, the moving direction and velocity keeps changing. The two kinds of walking patterns are intentional fast walking and slow unintentional walking. Fast walking occurs when the mouse wants to move from one position to another for a certain purpose such as eating and drinking. Slow walking occurs when the mouse roams in the cage and seems to have no idea on where to go and move. Figures 4(c) and 4(d) show the waveforms of acceleration and angular velocity of eight-second slow walking. The three-axis accelerations appear to change periodically. The small cycles in the readings may reflect the steps while the mouse is walking. However, the three-axis angular velocities show no repeated pattern but a dramatic change in x-axis. Rearing is usually found when the mouse is drinking or eating, because the water and food are placed on top of the cage. This action is called unsupported rearing, as the mouse’s front legs are unsupported. The other position is when the mouse rears along a wall and is called supported rearing, as the front legs touch the wall for support. This behavior starts and ends from the front legs lifting off the ground and touching back down. Capturing the complete behavior from the video is relatively routine. The variation of accelerations should be very large because the sensor orientation changes dramatically. Depending on the angle between the mouse and ground, the maximum variation of acceleration can reach 1g. Figures 4(e) and 4(f) show the acceleration and angular velocity of an eight-second complete rearing behavior. The three-axis acceleration undergoes changes and even overlaps. The rearing up and down actions can be easily found in the angular velocity waveforms, where the changing is most dramatic. Digging is determined when the mouse stores food or builds a comfortable place to stay. The mouse swings its front legs fast to throw the bedding particles around. At the same time, its head may dive into the bedding to check. However, the behavior is unpredictable because its ending cannot be clearly marked or observed. Figures 4(g) and 4(h) show the three-axis acceleration and angular velocity waveforms of a series of behaviors, including dig-stop-dig. The first digging lasts approximately 1.8 seconds, whereas the second one lasts approximately 1.6 seconds. During digging, both the acceleration and angular velocity change dramatically because the mouse’s front legs and head are both involved in this behavior. Shaking always happens while the mouse is sleeping. The mouse suddenly wakes up from sleeping and begins to shake intensely. The shaking gradually slows and stops, and then a new turn of shaking starts. The mouse usually performs several turns of shaking each time. This behavior is similar to sleeping, which is a long continuous behavior. However, shaking is much more intense in terms of collected sensor data than sleeping, as shown in Figures 4(i) and 4(j). The acceleration and angular velocity vary much during shaking than that while sleeping, and even close to that of walking. However, the orientation changes slightly while walking, but changes considerably while shaking.

Among all these five behaviors, sleeping, walking, digging, and shaking are all status-based behavior because they all have no apparent start or stop action. By contrast, rearing is a combination of movements that contain start, stop, and transition actions. The window size has limited influence on sleeping, walking, digging, and shaking, but has a critical influence on rearing. Therefore, the window size should be at least longer than the duration of rearing, the shortest duration of which is 2s from all the collected samples.

### 4.2 SVM Parameter Optimization

Two most important parameters of SVM are *gamma* and *C*. Intuitively, *gamma* defines a distribution after mapping training data into a new high dimension. A larger gamma means less support vectors, whereas a smaller gamma indicates more support vectors. The number of support vectors affects training and test speeds. *C* is a cost coefficient, which controls tolerance to error. Thus, *C* works as a threshold of decision function to accept or reject a training sample as a classification. A larger *C* means less error that the decision function tolerates, which causes an overfitting problem. A smaller *C* tolerates more errors, which leads to an underfitting problem.

A grid search with a five-folder cross validation is used to evaluate the influence of each pair of *gamma* and *C* on the classifier. Grid search tests all possible combinations of *gamma* and *C* in a given range. Figure 7 shows the validation accuracy of *gamma* (range: 0.1, 0.01, 0.001, 0.0001, 0.00001) and *C* (range: 0.1, 1, 10, 100, 1000), where an optimal combination of *gamma*=0.01 and *C*=10 can be found.

**Figure 7.**
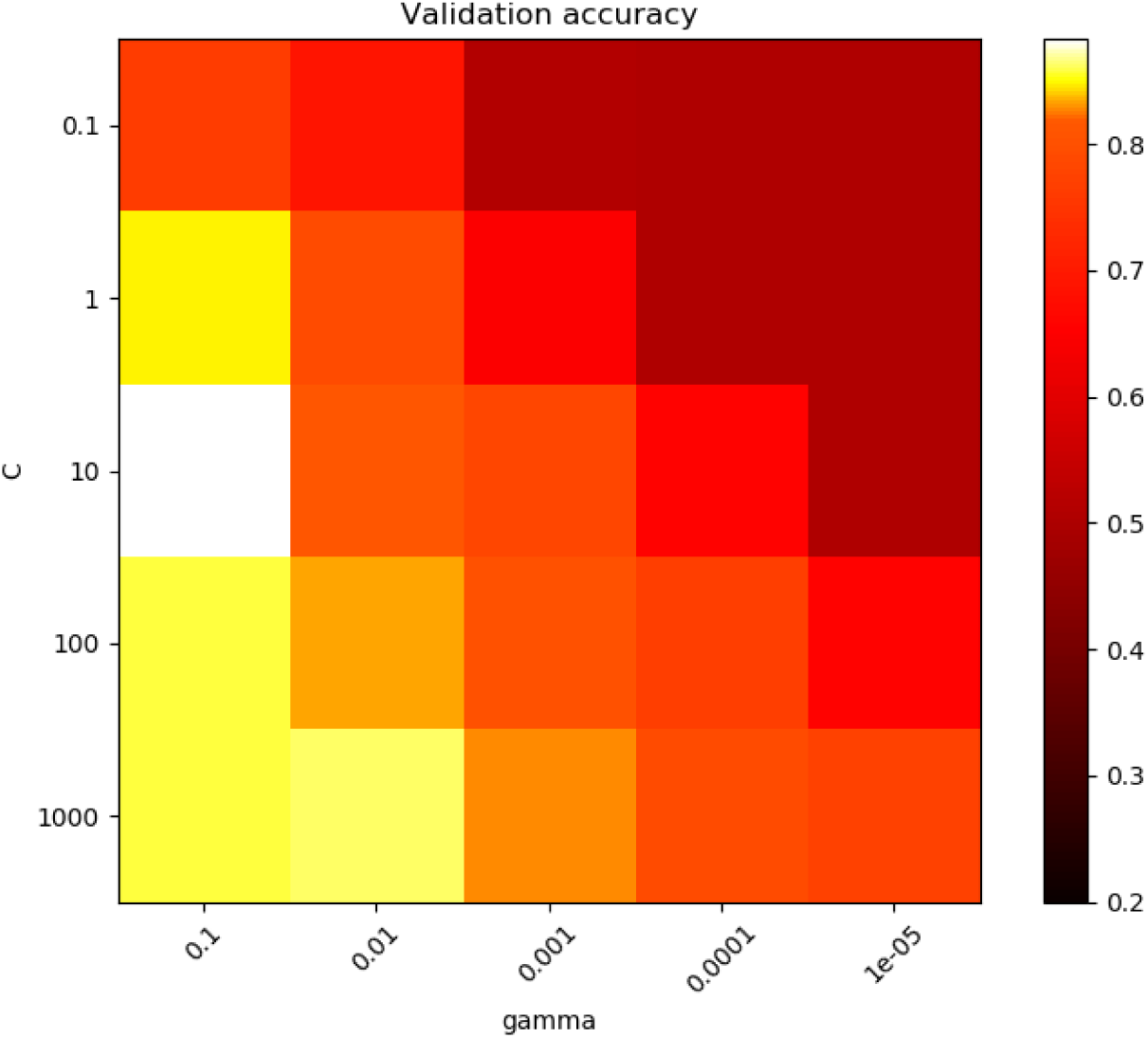
SVM parameter optimization

### 4.3 Feature Selection and Imbalanced Learning

As discussed in the preceding section, the SFS, SMOTE, and ClusterCentroids methods are used for feature and sample selection. A new balanced data set with a sample size *n* and feature size *m* can be easily generated by combining these two methods. Given that an infinite combination of the two parameters exists, *n*_*0*_ should be fixed. The optimal *m*_*1*_ is obtained and used in finding the optimal *n*_*1*_. The iteration stops at *m*_*k*_=*m*_*k-1*_ *or n*_*k*_=*n*_*k-1*_. Figure 8 shows the flow of the imbalanced learning. First, the imbalanced data set is split into training and test samples. The initial sample size *n*_*0*_ is set to the size of class with the fewest samples. Then, the following steps are performed:

**Figure 8.**
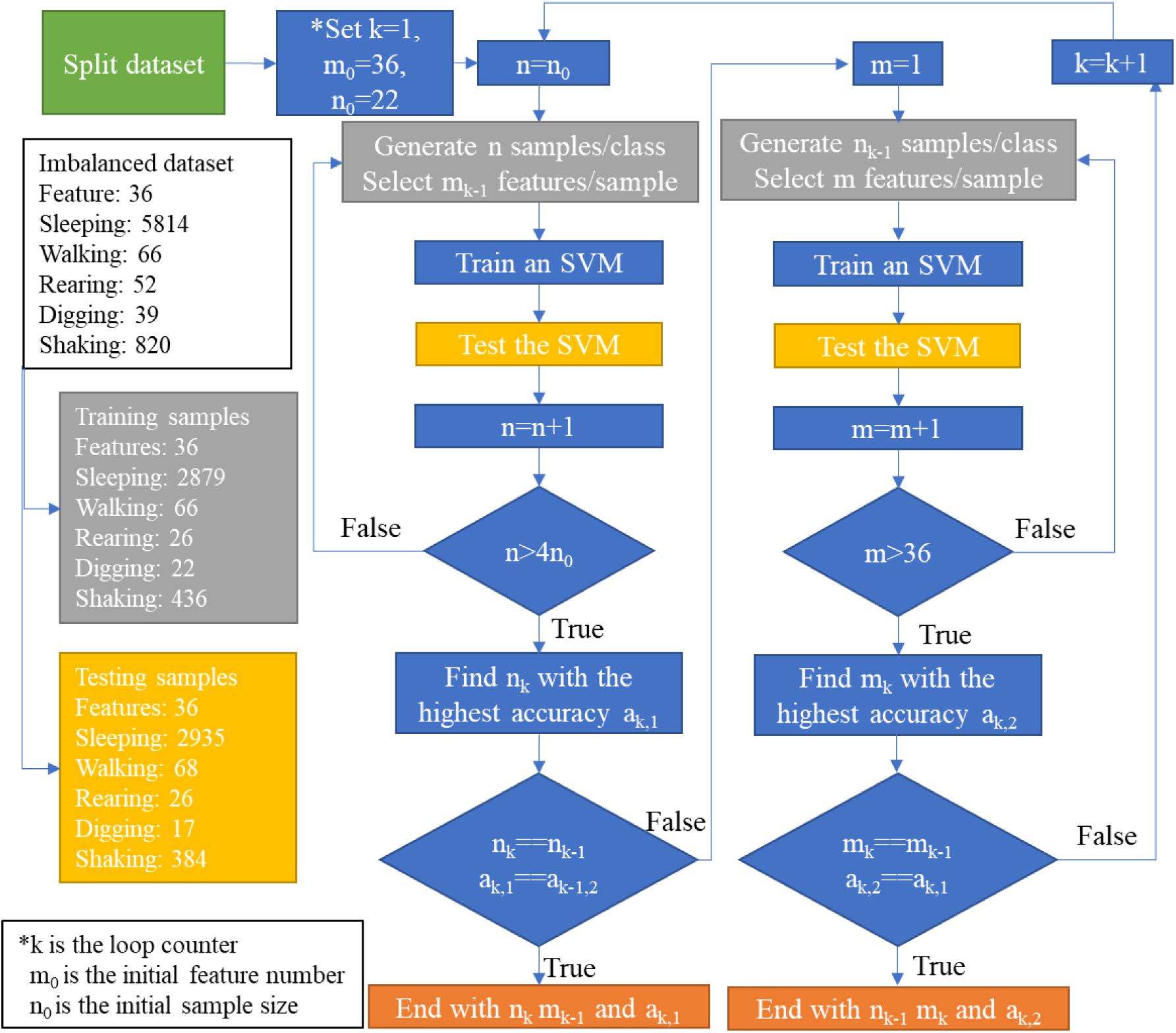
Flow chart of imbalanced learning

a. Find optimal sample size: Apply sample selection on the training samples with an initial size of six samples/class. Then, train an SVM using the generated data set with all 36 features/sample and derive the classification accuracy by testing the SVM with test samples. Increase the sample size from *n*_*0*_ to 4*n*_*0*_ with an increment of 1, and determine the highest average accuracy and corresponding sample size.
b. Find optimal feature number: Apply sample selection on the original training samples with updated sample size. Apply feature selection on the generated training dataset with initial number of 1 feature/sample to generate a new training dataset and train an SVM. Increase the feature number from 1 to 36 with an increment of 1, and determine the highest accuracy and corresponding feature number.

Optimization is achieved by the repetition of steps (a) and (b) until the two parameters (i.e., sample size and feature number) are covered. This process can be set automatically in the future. However, for purposes of illustration, an example is discussed as follows.

The procedure starts with an imbalanced dataset I and optimized SVM classifier. The average accuracy of the five behaviors is 48.07%. Sample selection is first applied to derive the current optimal dataset in the range of *n*_*0*_ to *4n*_*0*_ samples for each behavior with all 36 features. Figure 9(a) shows the results, where a highest average accuracy of 75.36% can be reached with an oversampling ratio of 3 for each behavior. Feature selection is then performed on the dataset with this oversampling ratio of 3, which is the optimal sample size found in the last step. Given that 36 features are present, the range of feature selection is from 1 to 36. Figure 9(b) shows the results, where a current optimal feature size of 33 is found with the highest average accuracy of 76.23%. Finally, 33-features data set proves that an oversampling ratio of 3 produced the best classifying results. The iteration stops at this point because feature and sample selections reached the same result. The optimized average accuracy reaches 76.23%. However, the optimal result for other windows sizes is not as good as that for a 2s window size. After implementing the optimization steps, the accuracy of minor classes including walking rearing and digging can be improved more than 58%, a considerable improvement, as shown in Table 4. But the accuracy of shaking drops 62.24% as the tradeoff. So, the total average accuracy increases 28.16%.

**Table 4.** Summary of the classification result in training imbalanced dataset I

| Behaviors | Sleeping | Walking | Rearing | Digging | Shaking | Average |
| --- | --- | --- | --- | --- | --- | --- |
| Testing sample | 2935 | 68 | 26 | 17 | 384 | 686 |
| Original accuracy | 99.18% | 16.18% | 3.80% | 23.53% | 97.66% | 48.07% |
| Optimized Accuracy<br>(with 3.05<br>oversampling ratio<br>and 33<br>features/samples) | 99.93% | 75% | 88.46% | 82.35% | 35.42% | 76.23% |
| Improvement | 0.75% | 58.82% | 84.66% | 58.82% | -62.24% | 28.16% |

**Figure 9.**
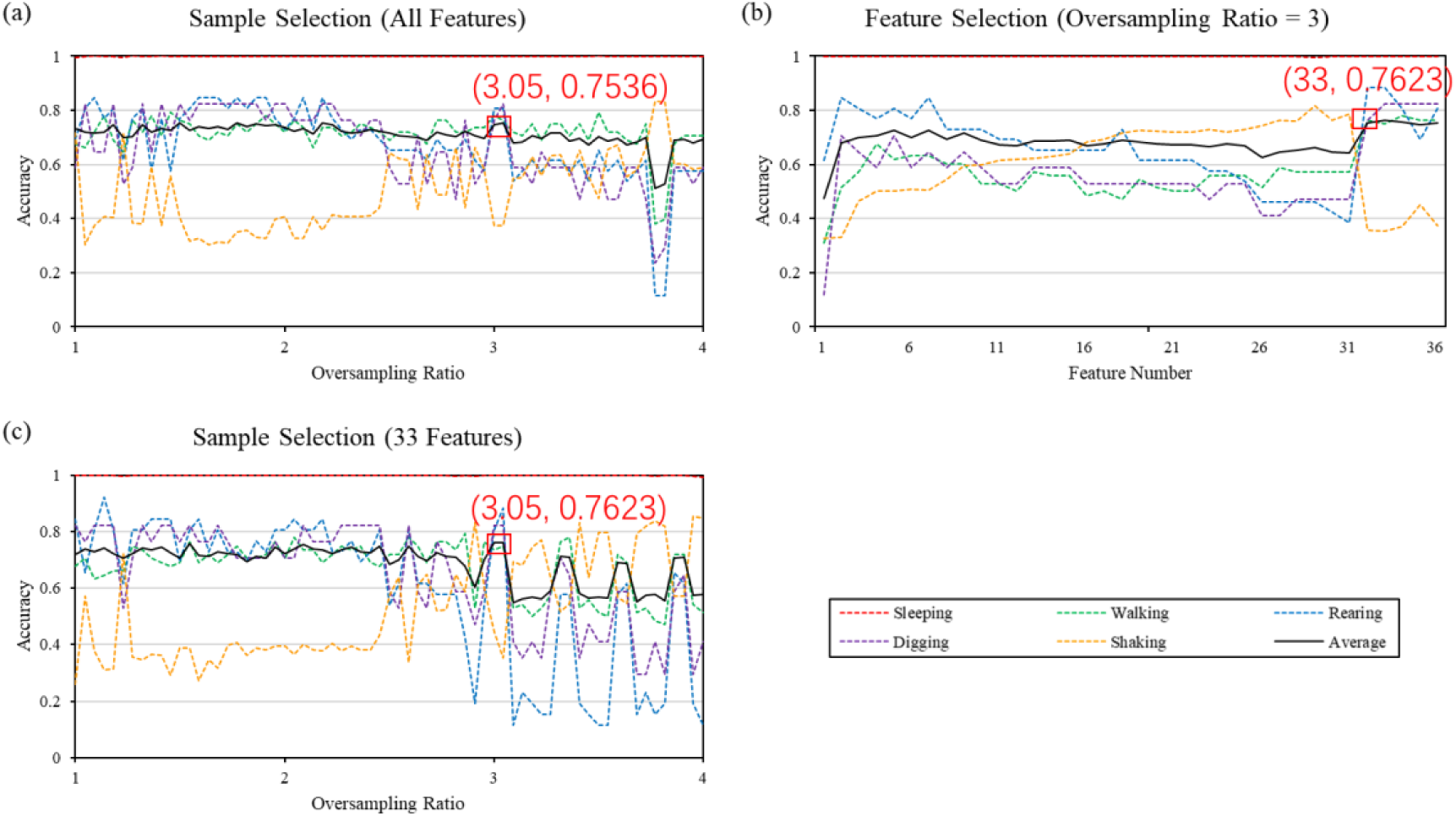
Classification results of each step in training imbalanced dataset I: from a to c

To prove the repeatability of the algorithm, a similar process is applied on imbalanced dataset II from another experiment. Table 5 compares the classification result before and after optimization, where the minor class shows a large accuracy improvement (48.68%). The optimization procedure stops at oversampling ratio of 3.34 and 18 features to obtain the highest average accuracy of 84.91%. However, the accuracy for walking and shaking decreases 9.42% and 5.13%. Finally, the average accuracy improves only by 7.31%.

**Table 5.** Summary of the classification result in training imbalanced dataset II

| Behaviors | Sleeping | Walking | Rearing | Shaking | Eating | Average |
| --- | --- | --- | --- | --- | --- | --- |
| Testing sample | 1095 | 138 | 76 | 604 | 299 | 442.4 |
| Original Accuracy | 92.42% | 94.93% | 42.11% | 77.81% | 81.94% | 77.84% |
| Optimized Accuracy<br>(with 3.34<br>oversampling ratio<br>and 18<br>features/sample) | 92.60% | 85.51% | 90.79% | 69.04% | 86.62% | 84.91% |
| Improvement | 4.75% | -9.42% | 48.68% | -5.13% | 4.68% | 7.31% |

According to the testing results of previous two imbalanced datasets, the accuracy of shaking drops after going through the optimization procedure. It seems the shaking samples have inverse relationship with other behaviors like walking, rearing, digging and eating. To validate this observation, shaking samples are removed from the two datasets and test other four behaviors with the proposed optimization procedure.

For imbalanced dataset I, the procedure also starts from sample selection with range from *n*_*0*_ to *4n*_*0*_ and all 36 features. Figure 10(a) shows the results that a highest average accuracy of 86.46% is found at an oversampling ratio of 2.2. The following feature selection proves that with the oversampling ratio of 2.2, the highest accuracy is achieved when the feature number is 36. The accuracies of walking and rearing are improved by the optimization procedure, as shown in table 6. After removing shaking, walking rearing and digging have 70.92% 67.03% and 46.47% accuracy increase, which proves that shaking interferes the classifier to correctly differentiate the mentioned three behaviors.

**Table 6.** Summary of the classification result in training imbalanced dataset I without shaking samples

| Behaviors | Sleeping | Walking | Rearing | Digging | Average |
| --- | --- | --- | --- | --- | --- |
| Testing sample | 2914 | 62 | 24 | 20 | 686 |
| Original accuracy | 99.73% | 87.1% | 70.83% | 70% | 81.91% |
| Optimized Accuracy | 99.90% | 96.77% | 79.17% | 70% | 86.46% |
| Improvement | 0.17% | 9.67% | 8.34% | 0% | 4.55% |

**Figure 10.**
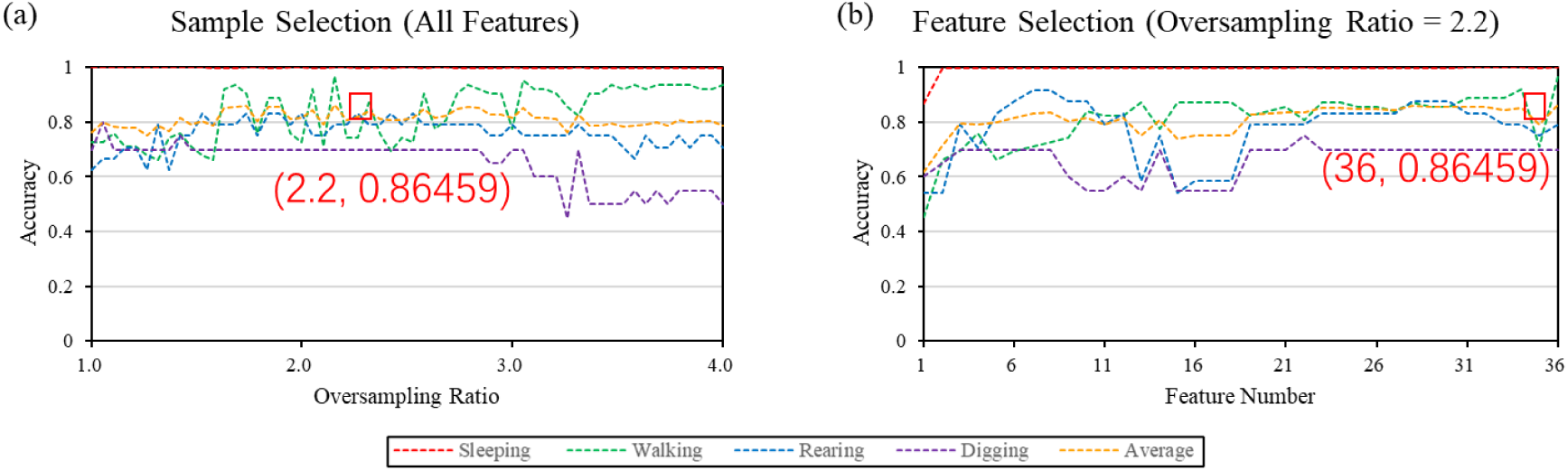
Classification results of each step in training dataset I without shaking behavior

Table 7 is the result of removing shaking samples from imbalanced dataset II. Compared with before removing shaking samples, the original accuracies of rearing and eating increased to 83.33% (from 42.11%) and 98% (from 81.94%). However, there is a tradeoff of 5.85% reduction in the accuracy of walking (i.e. from 94.93% to 89.68%, before optimization). Taken as a whole, removing shaking samples increases the overall average accuracy by 11.14% (i.e., from 84.91% to 94.37%, after optimization).

**Table 7.**
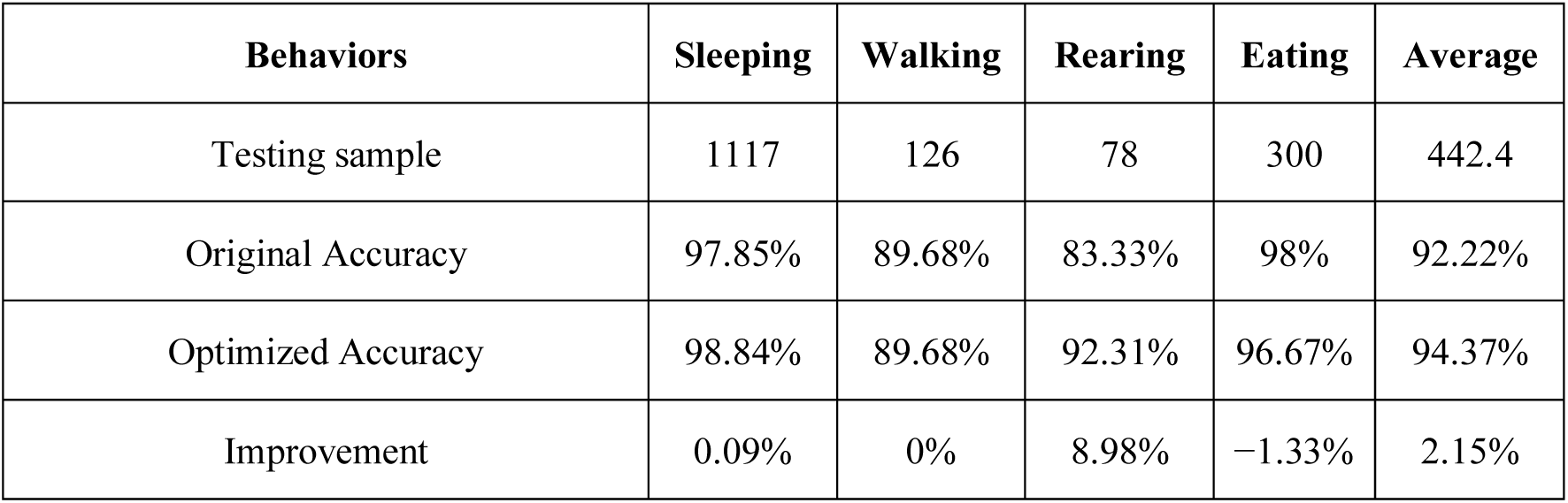
Summary of the classification result in training imbalanced dataset II without shaking samples

To further validate the result, imbalanced dataset III without shaking was used. Also, in dataset III, we included digging, grooming, drinking and scratching to see if our optimization algorithm works for less frequent behaviors, as compared to sleeping and walking. Table 8 shows the results before and after applying the optimization on the dataset. Consistent with the last dataset, the results show that, without the interference of shaking, the included five behaviors can be distinguished with over 90% accuracy, indicating that the classifier can classify these five behaviors with relative high performance.

**Table 8.** Summary of the classification result in training imbalanced dataset III

| Behaviors | Rearing | Digging | Grooming | Drinking | Scratching | Average |
| --- | --- | --- | --- | --- | --- | --- |
| Testing sample | 66 | 54 | 70 | 53 | 23 | 53.2 |
| Original Accuracy | 89.39% | 96.29% | 95.71% | 94.33% | 60.86% | 87.32% |
| Optimized Accuracy | 89.39% | 100% | 94.29% | 98.11% | 100% | 96.35% |
| Improvement | 0% | 1.86% | -2.85% | 3.78% | 34.79% | 8.73% |

## 5. Discussion

As mentioned in previous sections, fixed-time-window segmentation is used in the proposed data analysis methodology. Different behaviors, such as *sleeping, walking, rearing, digging, shaking, eating, drinking, grooming*, and *scratching*, are identified manually through the recorded video (usually two hours each). The start and end times of each identified behavior are marked down to extract corresponding motion data from the entire log file of the experiment. Thus, various motion data and corresponding behavior classes are obtained. Data are then segmented by a fixed time window into numerous samples for feature extraction. As discussed above in the behavior motion visualization, the window size does not affect several behaviors, but may have a minimum for other behaviors. A suitable time window could segment data and avoid breaking the behavior into pieces. In this study, each piece of motion data corresponded to one behavior label. However, a piece of raw data acquired from a mouse may contain more than one behavior. If a fixed-time window is used to segment it into samples, the split point might not lie on the boundary of two behaviors, which would create the situation where one sample contains more than one behavior. This sample cannot be correctly classified using a machine learning algorithm. For normal usage, the entire system can be used to recognize animal behavior by a trained model, and the data obtained from animals becomes unlabeled. Such problem can be solved using an adaptive-window-segmentation method, which can detect the difference between two connected behaviors and split them. Thus, the window size follows the duration of one behavior. Figure 11 shows the main steps of this method. A base and extended window is used to achieve the adaptive window segmentation. For a new segment, the initial length is the duration of a base window. Then, the correlation between the segment and the next extended window is calculated to decide if the two windows belong to the same behavior. If so, the two windows are merged, and the same operation is applied on next extended window to change the window size until all windows differ. Otherwise, the base window is an independent behavior and the operation stops.

**Figure 11.**
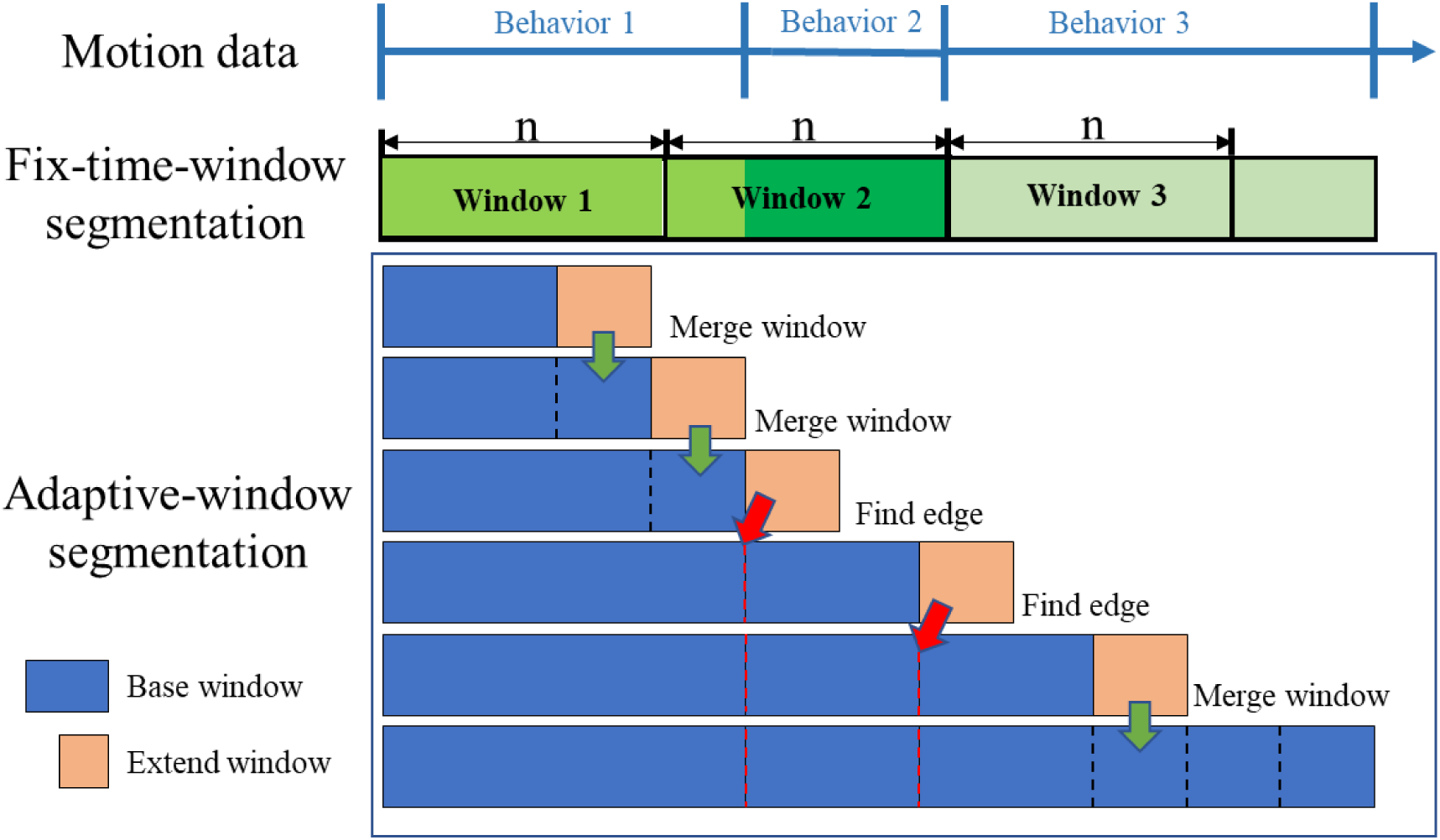
Comparison between fix-time-window and adaptive-window segmentations

The other problem is that the presence of shaking samples affected the classification accuracies of rearing and digging. Before optimization of dataset I, the accuracies of walking, rearing and digging are very low because of the relatively small sample sizes as compared with those of sleeping and shaking as shown in Table 4. During the optimization procedure as shown in Figure 12, the trends of the accuracies for walking, rearing and digging are totally opposite to that of shaking. This phenomenon is not only found in the oversampling ratio ranged from 1 to 4 (Figure 9c) but also appears in range beyond a ratio of 4 (Figure 12). The results of dataset II also indicate the contradiction between rearing and shaking samples’ accuracies. In other words, the proposed 36 features extracted from motion data may not be able to classify shaking from the others. There are two ways to solve this problem. First, more unique features must be identified. Second, time-domain signal processing methods such as Recurrent Neural Network (RNN) [43] is integrated to reveal the relationship of different states of one behavior. In this method, a continuous behavior will be split into pieces, which acts as the basic elements to form a behavior. Thus, different behaviors are different combinations of these elements. This method avoids the direct feature extraction of a single behavior as a whole.

**Figure 12.**
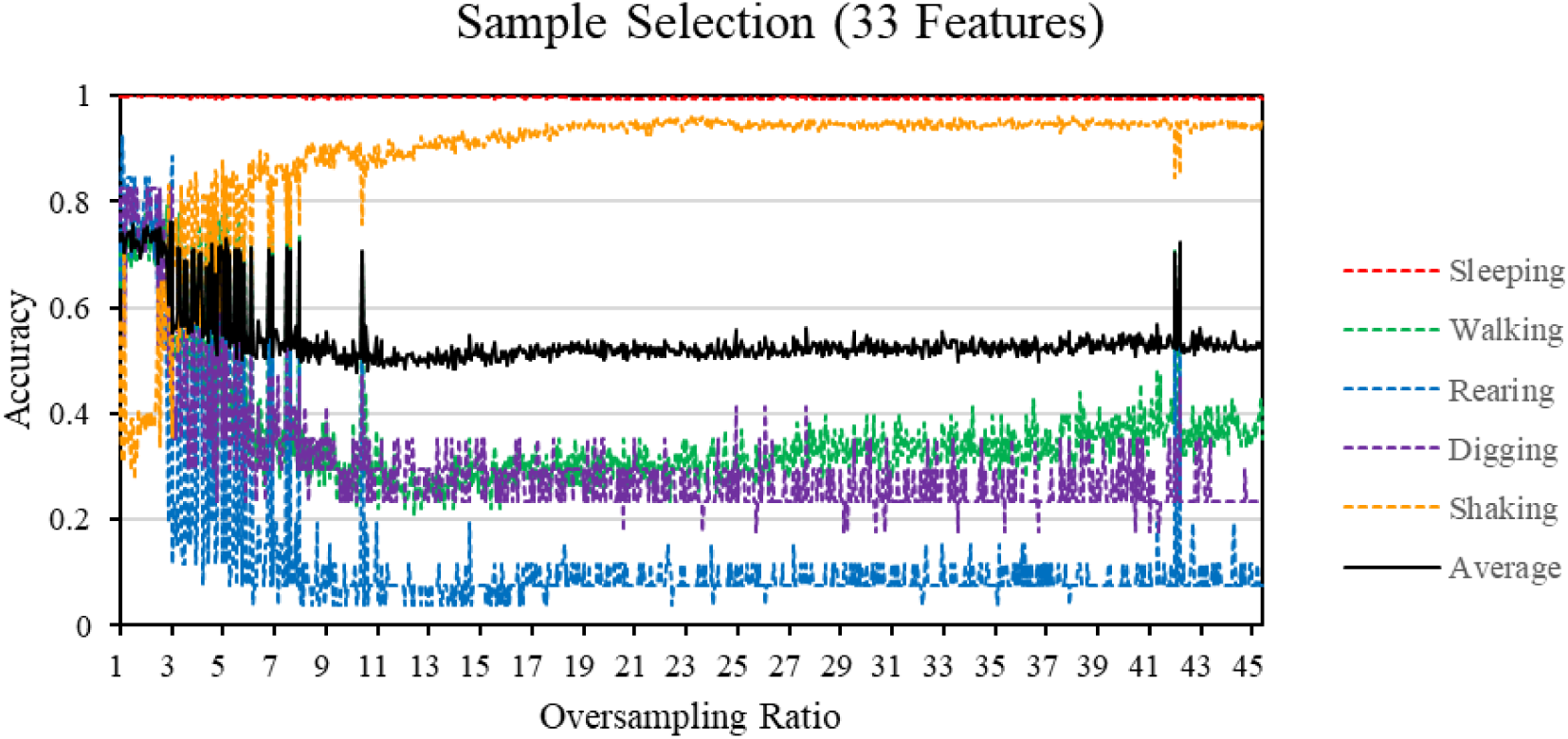
Extending the sample selection of imbalanced dataset I to higher oversampling ratio

## 6. Conclusion

In this study, a wireless AI-powered IoT sensors (AIIS) system for laboratory mice motion recognition utilizing embedded micro-inertial measurement unit for drug-testing applications is introduced. Animal motion data acquired by the AIIS are further classified into five behaviors, including *sleeping, walking, rearing, digging*, and *shaking*. The processing methods include data segmentation, feature extraction, feature selection, imbalanced learning, and machine learning. The acceleration and angular velocity waveforms of the five behaviors show the details of the motions. The result of using a tuned SVM to classify these five behaviors under imbalance distribution reaches 48.07% average accuracy. To improve the performance, a series of operations, including sample and feature selection, are applied to rebuild the training samples. Finally, the average accuracy of classifying five behaviors increases to 76.23% using the model trained by the rebuilt training samples. After removing *shaking* from these five behaviors, the average accuracy reaches 86.46% in classifying other four behaviors based on the rebuilt balanced training samples. For less frequent behaviors including *rearing, digging, grooming, drinking* and *scratching*, the average accuracy is 96.35%.

## Acknowledgements

This research was funded by the Hong Kong Innovation Technology Commission under Project UIM/382; and JLFS-RGC-Joint Laboratory Funding Scheme under Project JLFS/E-104/18

